# Structure, function and variants analysis of the androgen-regulated *TMPRSS2*, a drug target candidate for COVID-19 infection

**DOI:** 10.1101/2020.05.26.116608

**Authors:** Alessia David, Tarun Khanna, Melina Beykou, Gordon Hanna, Michael J E Sternberg

**Author notes:** Corresponding author*: Dr Alessia David, Centre for Integrative Systems Biology and Bioinformatics, Department of Life Sciences, Imperial College London, London SW7 2AZ, UK.

## Abstract

SARS-CoV-2 is a novel virus causing mainly respiratory, but also gastrointestinal symptoms. Elucidating the molecular processes underlying SARS-CoV-2 infection, and how the genetic background of an individual is responsible for the variability in clinical presentation and severity of COVID-19 is essential in understanding this disease.

Cell infection by the SARS-CoV-2 virus requires binding of its Spike (S) protein to the ACE2 cell surface protein and priming of the S by the serine protease TMPRSS2. One may expect that genetic variants leading to a defective TMPRSS2 protein can affect SARS-CoV-2 ability to infect cells. We used a range of bioinformatics methods to estimate the prevalence and pathogenicity of TMPRSS2 genetic variants in the human population, and assess whether TMPRSS2 and ACE2 are co-expressed in the intestine, similarly to what is observed in lungs.

We generated a 3D structural model of the TMPRSS2 extracellular domain using the prediction server Phyre and studied 378 naturally-occurring TMPRSS2 variants reported in the GnomAD database. One common variant, p.V160M (rs12329760), is predicted damaging by both SIFT and PolyPhen2 and has a MAF of 0.25. Valine 160 is a highly conserved residue within the SRCS domain. The SRCS is found in proteins involved in host defence, such as CD5 and CD6, but its role in TMPRSS2 remains unknown. 84 rare variants (53 missense and 31 leading to a prematurely truncated protein, cumulative minor allele frequency (MAF) 7.34×10−4) cause structural destabilization and possibly protein misfolding, and are also predicted damaging by SIFT and PolyPhen2 prediction tools. Moreover, we extracted gene expression data from the human protein atlas and showed that both ACE2 and TMPRSS2 are expressed in the small intestine, duodenum and colon, as well as the kidneys and gallbladder.

The implications of our study are that: i. TMPRSS2 variants, in particular p.V160M with a MAF of 0.25, should be investigated as a possible marker of disease severity and prognosis in COVID-19 and ii. in vitro validation of the co-expression of TMPRSS2 and ACE2 in gastro-intestinal is warranted.

## INTRODUCTION

The transmembrane protease serine type 2 (TMPRSS2) protein has a key role in severe acute respiratory syndrome (SARS)-like coronavirus (SARS-CoV-2) infection, as it is required to prime the virus’ spike (S) protein, thus facilitating its entry into target cells (Hoffmann et al., 2020), (Shang et al., 2020). TMPRSS2 is characterized by androgen receptor elements located upstream its transcription site (Lin et al., 1999). It is expressed in lung and bronchial cells (Lukassen et al., 2020), but also in the colon, stomach, pancreas, salivary glands, prostate and numerous other tissues (Vaarala et al., 2001). Moreover, TMPRSS2 has recently been shown to be co-expressed with the angiotensin-converting enzyme 2 (ACE2) – the SARS-CoV-2 cellular receptor (Zhou et al., 2020) (Hoffmann et al., 2020) – in bronchial and lung cells (Lukassen et al., 2020).

TMPRSS2 cleaves and activates the S protein of several coronaviruses, including the severe acute respiratory syndrome-related coronavirus (SARS-CoV) (Hoffmann et al., 2020) and the Middle East respiratory syndrome-related coronavirus (MERS-CoV) (Du et al., 2017), facilitating virus-cell membrane fusion and viral infection. TMPRSS2 is also crucial for the proteolytic cleavage and activation of hemagglutinin (HA) in the influenza A virus (Böttcher et al., 2006).

Studies in TMPRSS2-KO mice reported reduced SARS-CoV and MERS-CoV replication in lungs compared to wild-type mice, and a reduced proinflammatory viral response, particularly cytokine and chemokine response via the Toll-like receptor 3 pathway (Iwata-Yoshikawa et al., 2019). In vitro studies has shown that TMPRSS2 inhibitors prevent bronchial cell infection by SARS-CoV (Kawase et al., 2012). In animal studies, mice infected with SARS-CoV and treated with the serine protease inhibitor Camostat Mesilate showed a high survival rate (Zhou et al., 2015). Recently, Camostat Mesilate (which, in Japan, is already approved for use in patients with chronic pancreatitis and postoperative reflux esophagitis) was shown to block SARS-CoV-2 lung cell infection *in vitro* (Hoffmann et al., 2020). Based on these encouraging results, a clinical trial to study the effect of a TMPRSS2 inhibitor in COVID-19 patients is under way (https://clinicaltrials.gov/ct2/show/NCT04321096).

Since pharmacological block of TMPRSS2 prevents cell entry and infection by coronavirus, one could postulate that naturally occurring *TMPRSS2* genetic variants affecting the structure and function of the TMPRSS2 protein may confer some protection from SARS-CoV-2 infection. We performed a computational bioinformatics analysis on *TMPRSS2* variants reported in GnomAD, the database of population genetic variations, to identify variants that could result in TMPRSS2 loss of structure/function, and assess their prevalence in the general population using data from the GnomAD database. Moreover, we explored the co-expression of *TMPRSS2* and *ACE2* in extrapulmonary tissues using data from the Human protein Atlas, focusing on the intestine in view of the high prevalence of diarrhoea and gastrointestinal symptoms in COVID-19 patients (Chen et al., 2020).

## METHODS

*TMPRSS2* variants, with their global and gender-specific minor allele frequency (MAF), were extracted from GnomAD (Karczewski et al., 2017). Cumulative MAF was calculated as the sum of the MAFs for each variant.

In order to assess the structural impact of *TMPRSS2* variants, the three-dimensional structure of the TMPRSS2 was required. Currently, no experimental 3D structure for TMPRSS2 is available. We, therefore, generated a 3D structural model using our in-house Phyre homology modelling algorithm (Kelley et al., 2015). The FASTA sequence of TMPRSS2 was obtained from the UniProt protein knowledge database (The UniProt Consortium, 2017) (UniProt Id O15393, corresponding to 492 amino acid transcript Ensembl ID ENST00000332149.10). We deposited the 3D coordinates of the model in our PhyreRisk (Ofoegbu et al., 2019) database (http://phyrerisk.bc.ic.ac.uk/search?action=fresh-search&searchTerm=TMPRSS2). For completeness, SWISS-MODEL (Waterhouse et al., 2018) was also implemented to model TMPRSS2.

Model quality was assessed using: i. VoroMQA (Voronoi tessellation-based Model Quality Assessment) (Olechnovič and Venclovas, 2017), a statistical-based tool based on inter-atomic and solvent contact areas, which returns both a local and global score between 0 and 1. A global score > 0.4 indicates a good model and < 0.3 a poor model, models with >0.3 score <0.4 cannot be correctly classified; ii. QMEANDisCO (Studer et al., 2020), which uses distance restraints calculated from experimental structures of proteins homologous to the query protein and returns a plot of local IDDT (distance difference comparison) scores for each residue between 0 and 1. A global IDTT >0.6 indicates a good model structure. iii. ProSA (Wiederstein and Sippl, 2007). The latter returns: 1. the Z-score, which indicates the overall model quality of the target structure compared to the Z-score for all experimentally determined structures in PDB (X-rays and NMR); 2. the residue energy, which indicates the local model quality and is calculated over a 10- and 40-residue window. Negative energy values indicate good quality structure.

The impact of each variant on TMPRSS2 protein structure was assessed by analysing the following 16 features, using our in house algorithm Missense3D (Ittisoponpisan et al., 2019): breakage of a disulfide bond, hydrogen bond or salt bridge, introduction of a buried proline, clash, introduction of hydrophilic residue, introduction of a buried charged residue, charge switch in a buried residue, alteration in secondary structure, replacement of charged with uncharged buried residue, introduction of a disallowed phi/psi region, replacement of a buried glycine with any other residue, alteration in a cavity, replacement of cis proline, buried to exposed residue switch, replacement of a glycine located in a bend. In addition, we used the SIFT (Vaser et al., 2016) and Polyphen2 (Adzhubei et al., 2010) variant predictors, which mainly use evolutionary conservation to assess a variant’s effect.

The effect of variant rs12329760 was further assess using: i. CONDEL (González-Pérez and López-Bigas, 2011) which reports a weighted average of the scores from fatHMM and MutationAssessor, and ii. FoldX5 force field (Schymkowitz et al., 2005) which calculate the stability of a protein based on the estimation of its free energy. A ΔΔG > 0 kcal/mol (calculated as: destabilizing effect. ΔΔG= ΔG_mut_ − G_wt_) was predicted to have a destabilizing effect.

Expression data for *TMPRSS2* and *ACE2* were extracted from the Human Protein Atlas (HPA) (Uhlén et al., 2015): TMPRSS2 at https://www.proteinatlas.org/ENSG00000184012-TMPRSS2 and ACE2 at https://www.proteinatlas.org/ENSG00000130234-ACE2. Three sources of RNA data are integrated in the HPA database: i. HPA-generated RNA sequencing data, ii. RNA-seq data from the Genotype-Tissue Expression Project (GTEx), which uses post-mortem tissue samples, and iii. RNA-seq values from the Functional Annotation of Mammalian Genomes 5 (FANTOM5) project. HPA provides normalized expression (NX) values. In brief, to take into account the removal of non-coding transcripts, the Transcripts Per Million (TPM) values within each sample were adjusted to sum to one million. Thereafter, library size and composition differences were accounted for by applying a trimmed mean of M-values (TMM) normalisation to the sample TPM values. Then Pareto scaling was applied at gene level within each of the data sources. Batch effects were removed when the tissue data from the three source datasets was integrated. Where a tissue has multiple sub-tissues, the reported NX value was the maximum NX value across all the sub-tissues. A consensus transcript expression level was produced for each gene and tissue, by selecting the maximum NX value for this combination across the three data sources. Further details are available from the HPA website http://www.proteinatlas.org.

## RESULTS

### TMPRSS2 protein

TMPRSS2 is composed of a cytoplasmic region (residues 1-84), a transmembrane region (amino acids 85-105) and an extracellular region (residues 106-492). The latter is composed of three domains: the LDLR class A (residues 112-149), the scavenger receptor cysteine-rich domain 2 (SRCR-2) (residues 150-242) and the Peptidase S1 (residues 256-489), which contains the protease active site: residues H296, D345 and S441. Two potential glycosylation sites are present in positions 213 and 249. A cleavage site at residues 255-256 has been shown to allow shedding of the extracellular region of TMPRSS2 (Afar et al., 2001).

No known experimental structure for TMPRSS2 is currently available. We therefore generated a 3D model of the extracellular region residues 145-491 corresponding to domains SRCR and Peptidase S1 (Figure 1 and Figure 2 and Table 1) using the transmembrane serine protease hepsin as a template (PDB: 1Z8G, chain A, X-ray structure with 1.55Å resolution. Model confidence 100%, sequence Id= 35% to target sequence; Model assessment:.VoroMQA score = 0.462, QMEANDisCO score =0.67, and ProSA Z score = −8.96, also see Supplementary Figure S1. RMSD Phyre model vs SWISS-MODEL =0.30Å). A second model covering residues 111-150 (LDLRA domain) was generated using as a template PDB: 4U8U (giant haemoglobin), chain O, resolution 3.2Å (model confidence 97%, sequence Id= 30% to target sequence. (ProSA Z score= −3.08. VoroMQA score= 0.267, however this may not accurately reflect model quality since the software was trained on chains with more than 99 residues). Given the similarity of the Phyre and the SWISS-MODEL structures, the subsequent analysis should not be particularly sensitive to which model was selected.

**Table 1.**
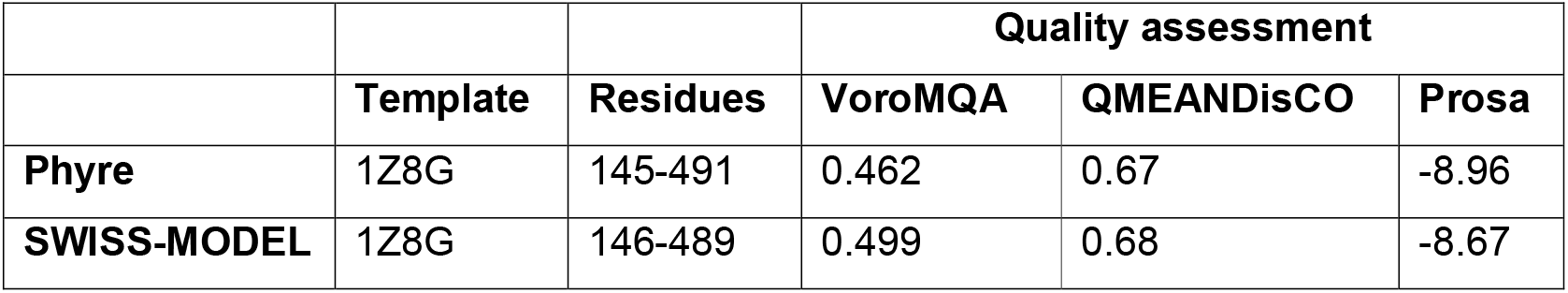
3D model of the SRCR and Peptidase S1 domain: template and quality assessment. Scores indicative of a good quality model are: VoroMQA >0.4, QMEANDisCO >0.6. For Prosa, the Z score ranges between −12 and −2.5 for X-ray structures of similar amino acid length (see suppl material).

**Figure 1.**
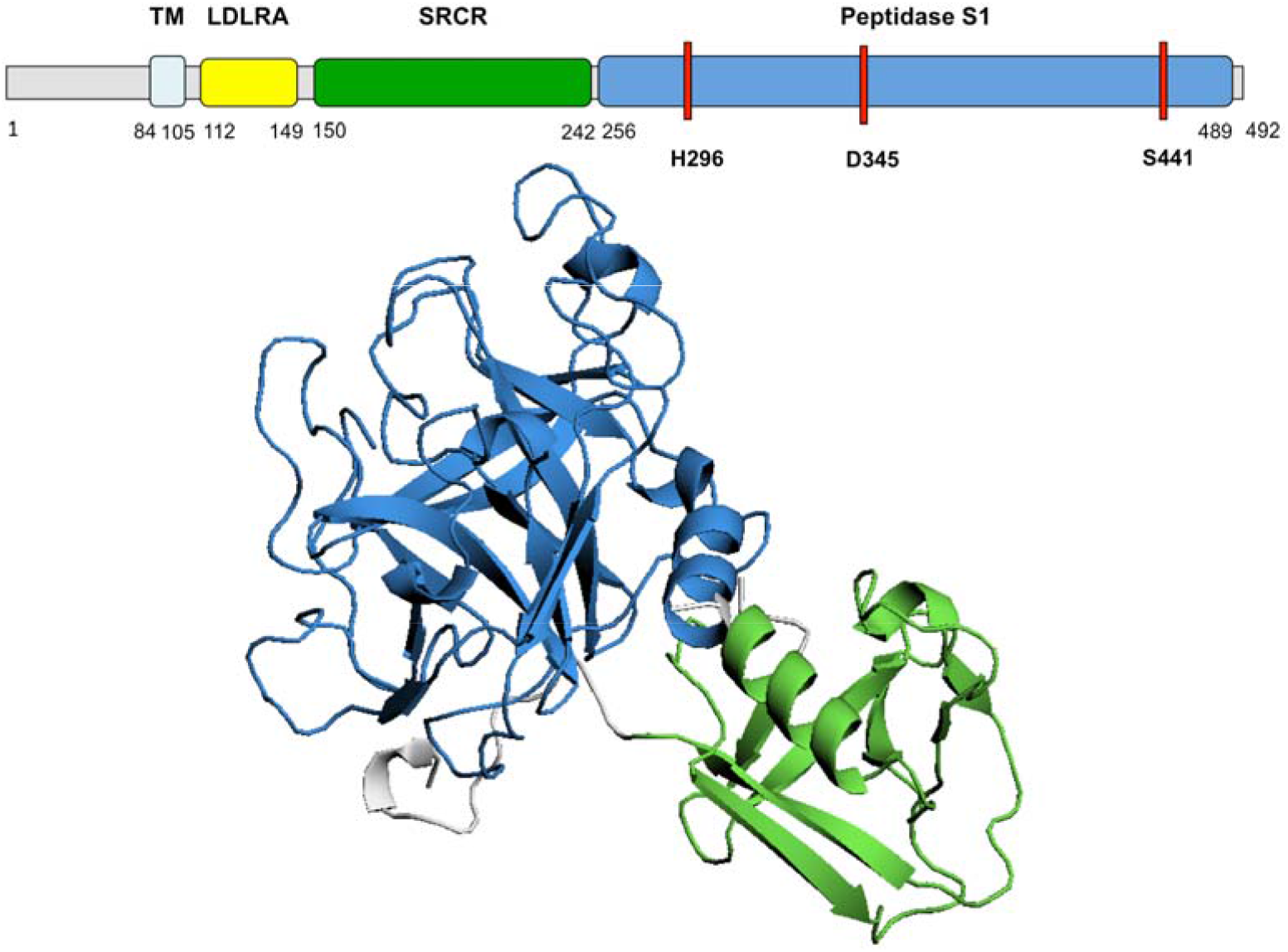
TMPRSS2 predicted 3D structure. Diagram of TMPRSS2 amino acid sequence and domains. The 3D model of TMPRSS2 domains SRCR and Peptidase S1 is presented. The active site, residues H296, D345 and S441, is highlighted in red on the amino acid sequence. TM, transmembrane domain; LDLRA, LDL-receptor class A; SRCR, scavenger receptor cysteine-rich domain 2; Peptidase S1, Serine peptidase.

**Figure 2.**
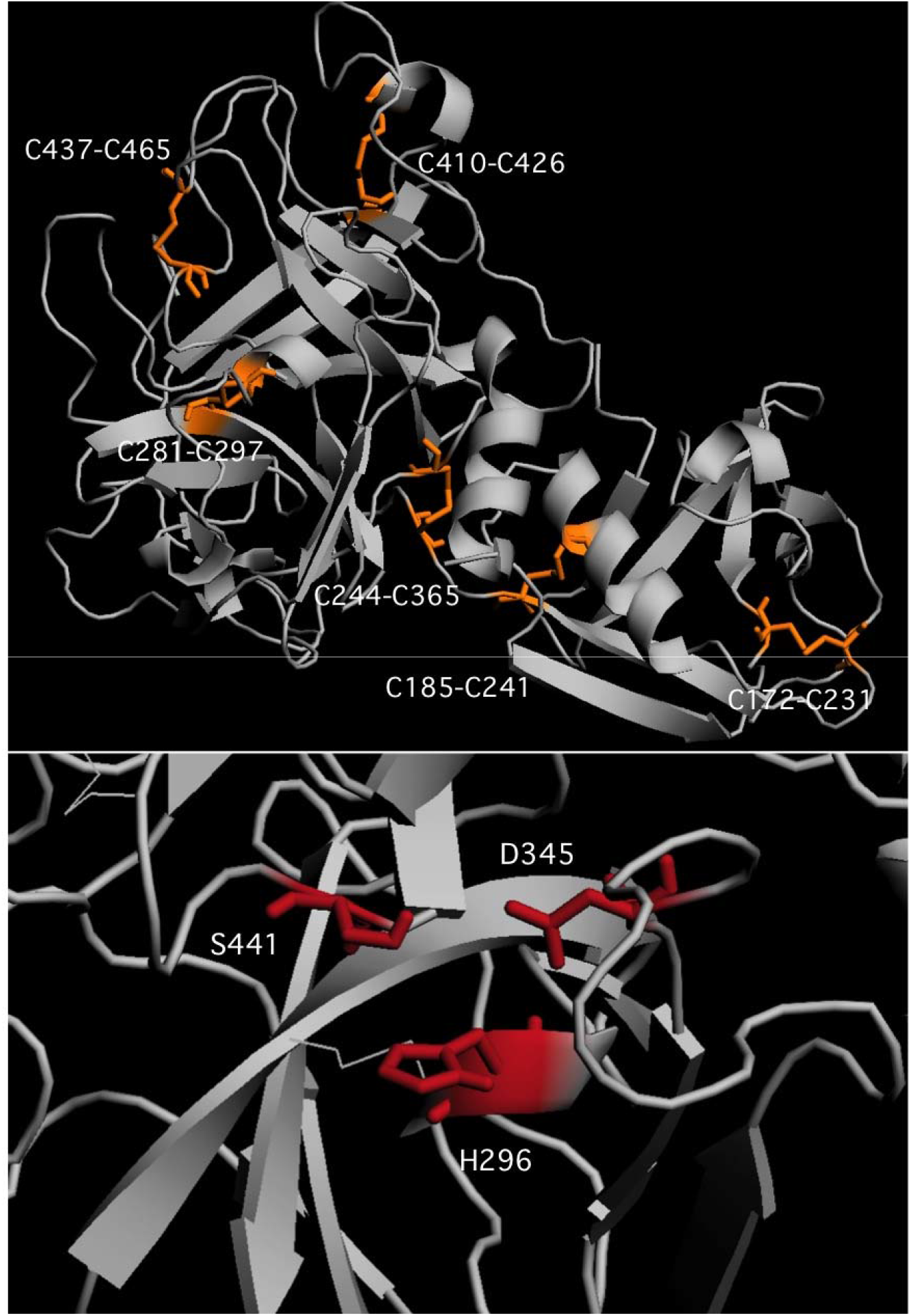
TMPRSS2 structure. Top panel: cysteine bonds in the SRCR and Peptidase S1 are presented in orange. Bottom panel: residues H296, D345 and S441 forming TMPRSS2 active site are presented in red

### TMPRSS2 variants

We analysed 378 naturally occurring *TMPRSS2* variants reported in the GnomAD database. One variant p.V160M (rs12329760) has a MAF of 0.248 in the population (0.2496 in males and 0.2488 in females), with 6.7% of individuals homozygotes for this variant (9,587 homozygotes out of 141,456 individuals sequenced as part of the GnomAd project). This variant is predicted damaging by both SIFT (score=0.01) and PolyPhen2 (score=0.937). CONDEL also predicts this substitution to be damaging (score=0.788). Valine 160 is a highly conserved, small hydrophobic residue within the SRCS domain and may have a structural or functional role within TMPRSS2 (Figure 3). On manual inspection, substitution of valine with the large methionine may cause some degree of steric clash within the structure. However, this was not supported by the change in free energy (ΔΔ*G*_FoldX_ −0.47 kcal/mol).

**Figure 3.**
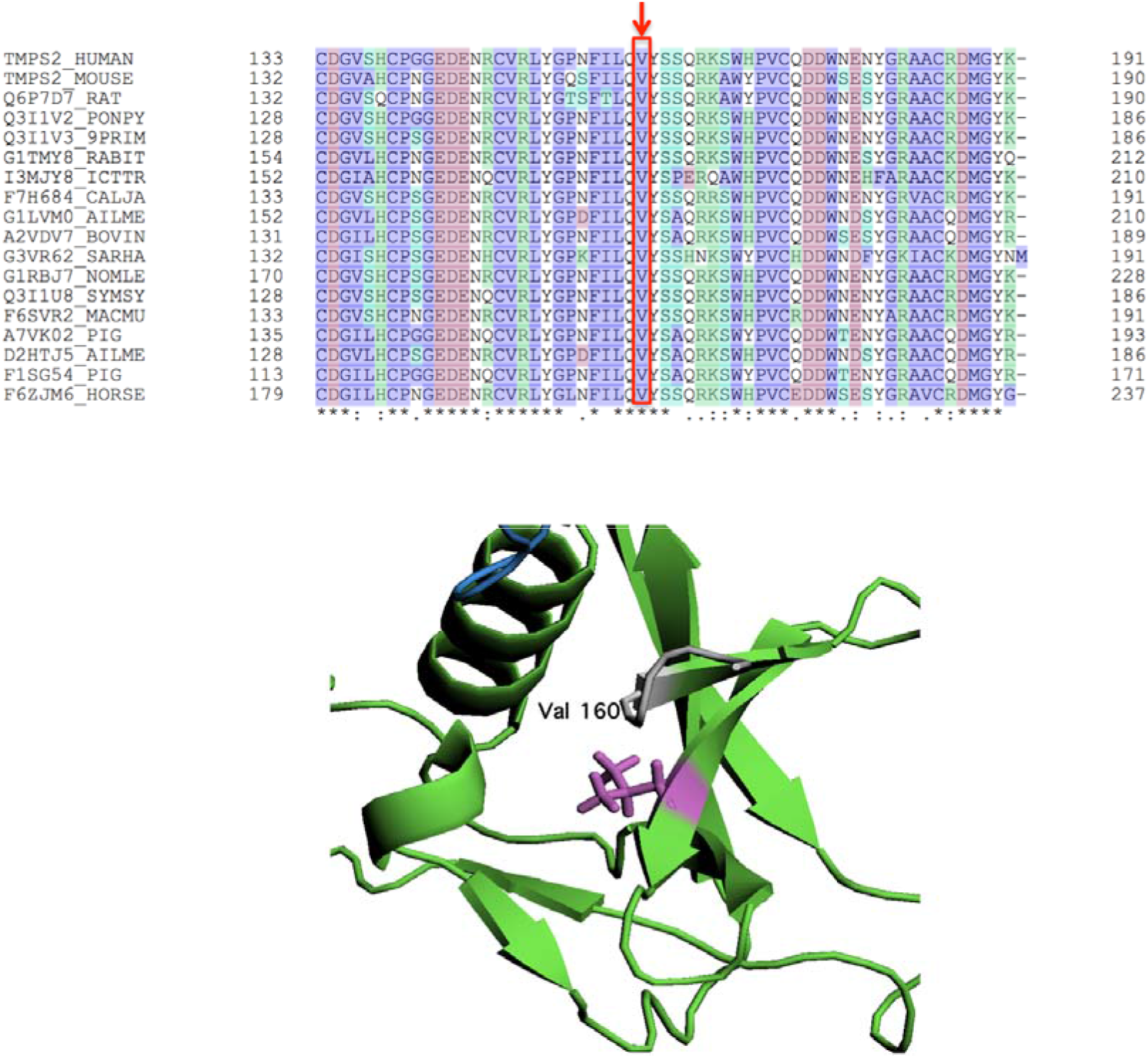
V160M genetic variant (rs12329760) in TMPRSS2 SRCR domain. The position of Valine 160 is presented within a multi sequence alignment and highlighted in magenta on the 3D structure of the SCRC domain

The p.V160M substitution may affect TMPRSS2 function. The SRCS is a highly conserved domain, whose function is still not fully understood, although a role in ligand and/or protein interaction has been proposed (Aruffo et al., 1997). Interestingly, this domain is present in several proteins involved in host defence, such as CD5, CD6 and Complement factor I (Freeman et al., 1990), (Resnick et al., 1994).

31 variants lead to a prematurely truncated protein (17 frameshift variants causing premature termination and 14 stop gain variants). 304 out of 334 missense variants were mapped onto the 3D structure and 62 variants were predicted structurally damaging by Missense3D. Of these, 12 variants (p.C113R, p.C113Y, p.C139R, p.C185Y, p.C244F, p.231S, p.C244R, p.C281F, p.C281R, p.C297Y, p.C410Y, p.C465Y) result in substitution of the invariable cysteine forming disulfide bonds and are highly likely to cause destabilization and possibly misfolding of the TMPRSS2 protein structure. Further analysis showed that two variants disrupt TMPRSS2 function: p.R255S (rs769655195), which abolishes the TMPRSS2 cleavage site and p.S441G (rs1292701415), which abolishes TMPRSS2 active site. However, both variants are extremely rare in the population (MAF<1×10−^5^) and are unlikely to be useful as a marker of SARS-CoV-2 infection severity.

167 variants were predicted damaging by SIFT and 152 by Polyphen2. 137 variants were predicted damaging by both SIFT and PolyPhen2, and of these, 53 variants were also predicted to cause structural damage, thus reinforcing the damaging effect predicted for these variants.

The 84 variants (53 missense and 31 leading to a prematurely truncated protein) that are predicted to cause loss of function with a high degree of confidence (predicted damaging by all three methods) are rare in the population. Their average MAF in the population is 9.67×10^−6^ and their cumulative MAF of 7.34×10^−4^, therefore unlikely to be helpful as a marker of SARS-CoV-2 infection severity in the general population.

### Both *TMPRSS2* and *ACE2* are expressed in the intestine

We extracted data on *TMPRSS2* and *ACE2* tissue expression from the Human Protein Atlas. As shown in Figure 4, both proteins are expressed in the gut: colon, small intestine and duodenum. Other sites of expression are the kidneys and gallbladder. Interestingly, although *TMPRSS2* and *ACE2* have recently been shown to be co-expressed in the lung and bronchial cells, *ACE2* lung expression is almost negligible in the HPA, thus highlighting that specific *in vitro* experiments are necessary to study the co-expression of *TMPRSS2* and *ACE2* in extrapulmonary tissues. Moreover, although expression of ACE2 arterial and venous endothelial cells and arterial smooth muscle cells was previously demonstrated (Hamming et al., 2004), it is not reported in the HPA.

**Figure 4.**
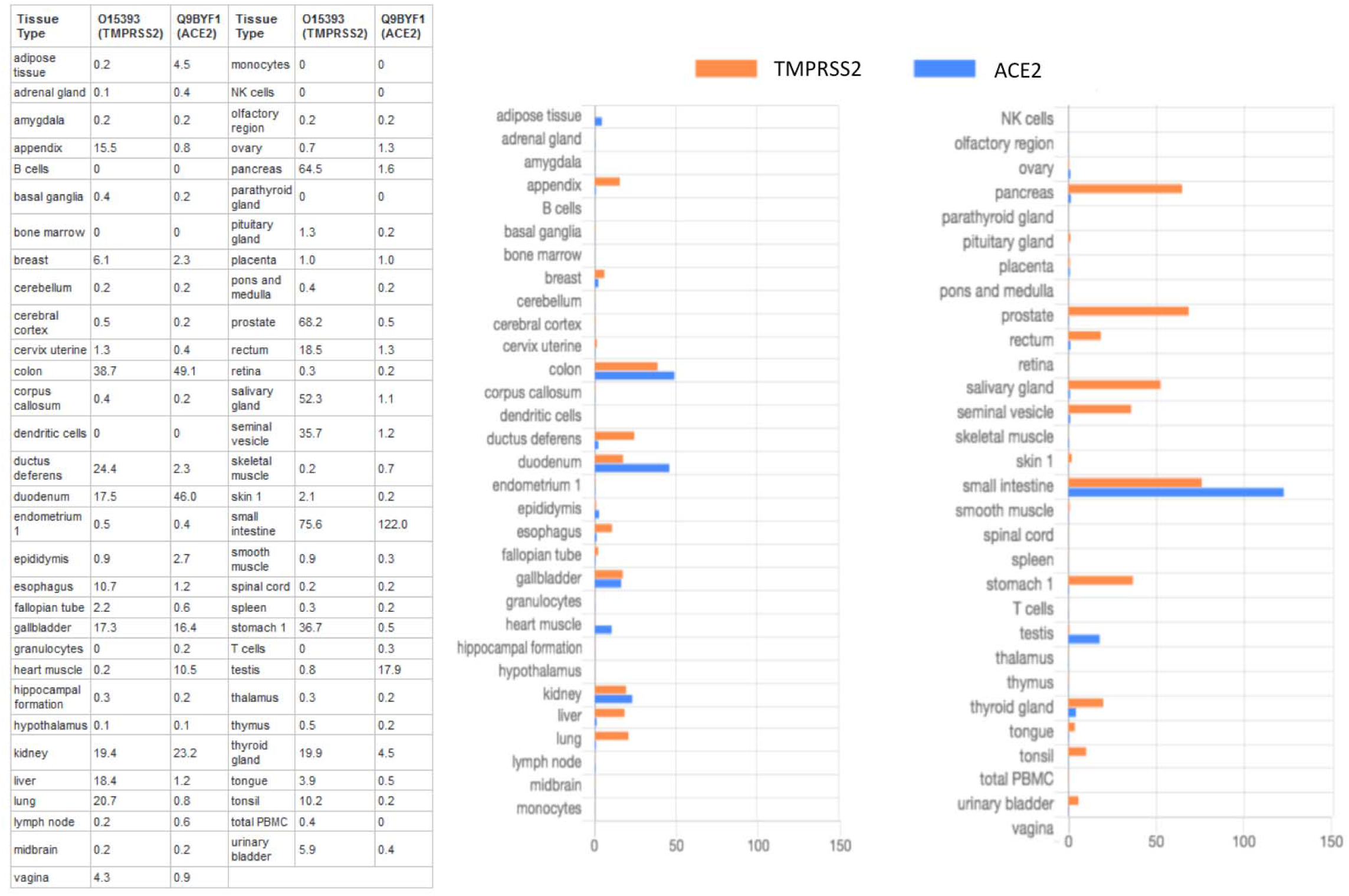
*TMPRSS2* and *ACE2* tissue expression.

## DISCUSSION AND CONCLUSION

As of May 2020, SARS-CoV-2 has infected millions of people around the world and has caused over 300,000 deaths. However, the true number of infected individuals remains unknown, as studies have suggested that many remain asymptomatic or have a mild symptoms (Day, 2020), (Song et al.). Naturally occurring genetic variations that result in a defective TMPRSS2 may explain why some individuals with COVID-19 develop mild disease. Our *in silico* analysis of *TMPRSS2* human variants shows that the predicted damaging substitution from valine to methionine at position 160 is a common genetic variant, present in almost 25% percent of the human population, with approximately 7% homozygotes according to GnomAd. *TMPRSS2* variants should be investigated further to understand the impact of a person’s genetic background on their clinical presentation and prognosis when contracting SARS-CoV-2 ,

Once SARS-CoV-2 infects an individual, it binds to ACE2 receptors on the surface of target cells. However, for the virus to enter the cells, double cleavage of the viral spike protein at the S1/S2 cleavage site and, subsequently, at the S2’ site is required. This allows viral fusion with the cell membrane and internalization (Hoffmann et al., 2020). TMPRSS2 is one the main cell surface proteases involved in the process of spike protein priming, although additional proteases, such as Furin and lysosomal cathepsin, are thought to be involved (Shang et al., 2020).

At present no known disease-association for *TMPRSS2* variants is known. A chromosomal aberration leading to gene fusion of *TMPRSS2* and its androgen promoter to *ERG*, known as the TMPRSS2-ERG fusion gene, has been identified in approximately one half of patients with prostate cancer (Haffner et al., 2010). Genetic variants leading to a defective and non-functioning *TMPRSS2* and *ACE2* are promising prognostic candidates for COVID-19 infection. Recent studies show that *ACE2* genetic variation is very rare in the population (Stawiski et al.) (MacGowan and Barton), thus making it an unlikely candidate to explain the wide range of symptoms (from asymptomatic to severely affected) observed in COVID-19 patients. We, therefore, focused on *TMPRSS2*, which together with *ACE2* plays an important role in SARS-CoV-2 infection. Although the majority of *TMPRSS2* variants are rare, the common variant V160M, which is predicted damaging, is an intriguing candidate for further study. Valine 160 is a highly conserved residue in the SRCS domain. It has been proposed that the latter may be involved in protein and/or ligand binding (Aruffo et al., 1997), however no known protein interaction partner is known to-date. The SRCR domain is often found in proteins involved in host defence (Freeman et al., 1990), (Resnick et al., 1994), thus raising the possibility that the role of TMPRSS2 in SARS-CoV-2 infection may extend beyond its peptidase activity related to viral protein priming.

Although respiratory tract infection is the most common manifestation of COVID-19, diarrhoea and gastrointestinal symptoms are also common. The co-expression of *ACE2* and *TMPRSS2* has been studied and experimentally ascertained in bronchial and lung cells, however co-expression of these proteins in the gut has not yet been investigated. We, therefore, used RNA expression data from the Human Protein Atlas (Uhlén et al., 2015) to show that *ACE2* and *TMPRSS2* are both expressed in the intestine. This suggests that, similarly to the lung, cells in the intestine may be susceptible to SARS-CoV-2 infection, resulting in gastro-intestinal symptoms in many patients, including children presenting with Kawasaki-like syndrome. This appealing pathogenetic hypothesis needs further *ad hoc in vitro* confirmation. It is, in fact, notable that the lung expression of *ACE2,* confirmed by dedicated *in vitro* studies (Lukassen et al., 2020), and its expression in arterial and venous endothelial cells and arterial smooth muscle cells (Hamming et al., 2004) are not reported in the HPA, thus suggesting that HPA data are incomplete.

In conclusion, SARS-CoV-2 is a new virus and still very little is known about the pathogenesis of COVID-19. Elucidating the molecular processes underlying SARS-CoV-2 infection and the factors responsible for the broad spectrum of disease severity observed in COVID-19 patients is crucial. Our data suggest that (i) *TMPRSS2* variants, particularly p.V160M which has a MAF of 0.25, should be investigated as a marker of disease prognosis and (ii) *in vitro* validation of the co-expression of *TMPRSS2* and *ACE2* in gastrointestinal cells is needed.

## Notes

The Authors declare no competing interest

### Competing Interest Statement

The authors have declared no competing interest.

