## Supplementary material for "Structure, function and variants analysis of the androgen-regulated *TMPRSS2*, a drug target candidate for COVID-19 infection": Figure S1

**Figure S1. Model analysis using PROSA for the 3D model covering domains SRCR and Peptidase S1 generated using Phyre and SWISS-MODEL.**

A) Overall model quality; B) local model quality.

### Phyre

A B

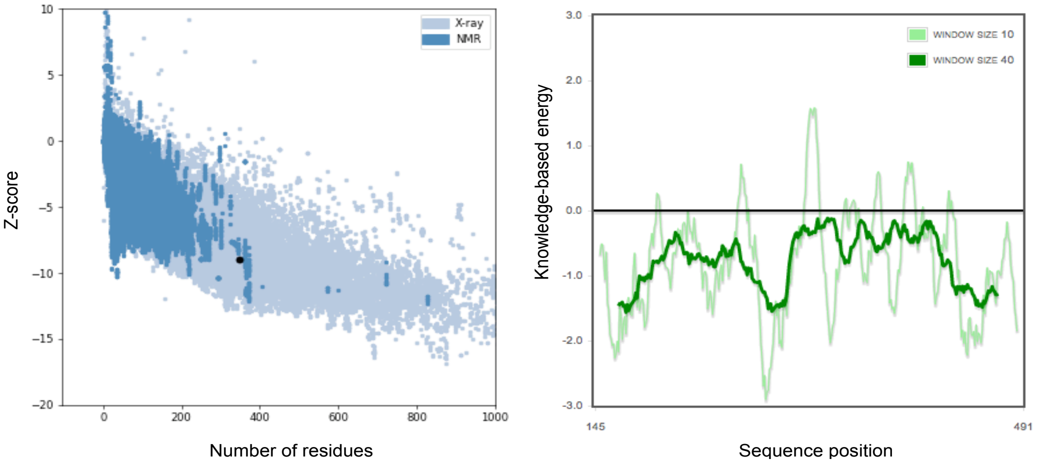


### SWISS-MODEL

A B

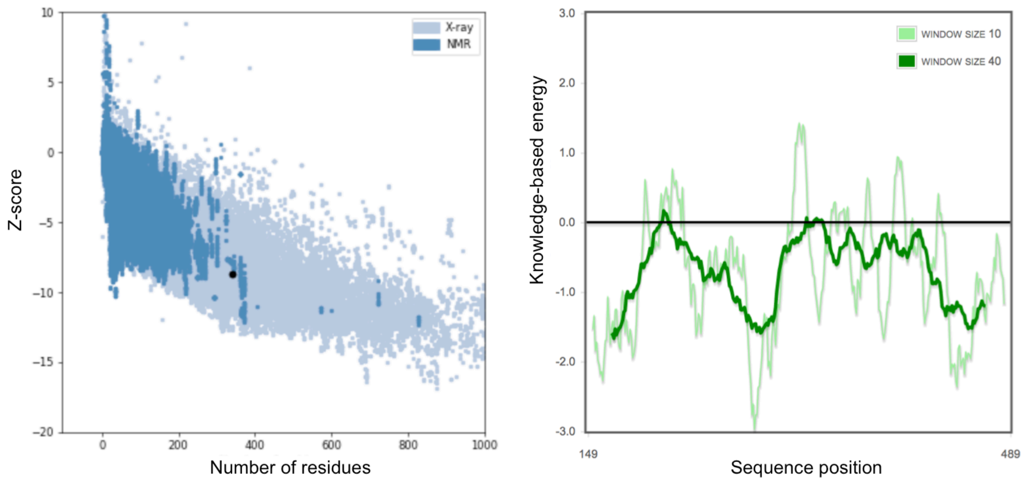
